# Environmental Noise Alters Neural Regulation Without Behavioral Impairment: A Pilot EEG Study

**DOI:** 10.1101/2025.11.04.685163

**Authors:** Egzona Morina

## Abstract

Environmental noise is a pervasive stressor that impairs attention and increases arousal, whereas natural soundscapes are linked to restoration and improved well-being. This pilot study tested whether exposure to natural auditory environments can buffer the neural strain induced by noise. Twelve healthy adults completed five cognitive tasks (DotProbe, Stroop, GoNoGo, NBack, Visual Search) under three auditory conditions: Noise (urban traffic), Nature (ambient natural sounds), and Control (silence with earplugs), while 32-channel EEG recorded ongoing activity. Behavioral accuracy remained high across all tasks (89.9 to 99.5 %), differing only for the Nback, which improved under Nature relative to Noise and Control. In contrast, EEG revealed robust environment-dependent modulation. Mixed-effects models (FDR-corrected *p* < .05) identified 17 significant condition effects across frequency bands and cortical regions. Nature increased baseline-corrected *δ*, *β*, and *γ* power, elevated *α/β* ratios, and enhanced the Engagement Index in posterior networks, all signatures of relaxed yet alert cortical states. Noise, by contrast, amplified *δ* and *θ* activity and raised *θ/α* ratios in parieto-occipital regions, indicating higher cognitive load and compensatory effort. Reliability analyses confirmed moderate within-condition stability of EEG measures (mean ICC = 0.53 to 0.61) and higher consistency for ratio-based indices. Together, these findings reveal that natural soundscapes promote efficient, low-effort neural organization, whereas urban noise elicits energetically costly activation despite preserved behavioral performance. The results establish electrophysiological markers of environmental stress and restoration, supporting biophilic design strategies for healthier and more cognitively sustainable work environments.

## 1 Introduction

Environmental exposures such as air pollution, heat, and noise increasingly threaten cognitive performance and well-being (Block and Calderon-Garciduenas, 2009; Bhui et al., 2023; Xiang et al., 2025; Arjunan and Rajan, 2020). While cardiopulmonary impacts of these stressors are well documented (Alahmad et al., 2023), far fewer studies have examined their influence on brain function. Emerging evidence links fine particulate matter (PM_2.5_), volatile organic compounds (VOCs), and ambient heat to neuroinflammation, altered neurotransmission, and structural brain changes (Calderon-Garciduenas et al., 2015; Costa et al., 2017; Beyer et al., 2024). Chronic noise exposure, in particular, has been associated with reduced working-memory capacity, impaired executive control, and heightened stress physiology (Hahad et al., 2020; Guo et al., 2017). These cognitive and biological effects are increasingly relevant in modern workplaces, where mental fatigue and distraction degrade creativity, decision-making, and general well-being. Theoretical frameworks such as Attention Restoration Theory (ART) propose that contact with natural environments replenishes depleted attentional resources and facilitates recovery from cognitive fatigue (Kaplan, 1995; Berman et al., 2008). Empirical work supports this view: visual or auditory exposure to nature reduces physiological arousal, improves task engagement, and enhances mood (Liu et al., 2024; Zhang et al., 2025). Conversely, even moderate background noise can elevate catecholamine stress markers and reduce intrinsic task motivation, suggesting that environmental noise constitutes a subtle but chronic cognitive stressor (**?**Evans and Johnson, 2000; Babisch et al., 2001; Sakellaris et al., 2016; Passchier-Vermeer and Passchier, 2000; Chen et al., 2025). Recent neuroimaging studies reinforce these behavioral findings. Forest soundscapes enhance attention-network connectivity and working-memory performance relative to urban noise (Stobbe et al., 2024). Together, this evidence indicates that natural environments promote focused and efficient cognitive states, whereas noisy or unpredictable ones disrupt attentional regulation. Despite this progress, neuroscientific understanding of how environmental context shapes moment-to-moment brain regulation remains limited. Most studies either rely on large-scale epidemiological data linking regional exposure indices to cognitive decline, or on highly constrained laboratory paradigms with limited ecological validity. Few investigations have combined real-time EEG recording with controlled manipulations of environmental soundscapes in realistic occupational settings. As a result, the neural dynamics underlying environmental stress and restoration are still poorly characterized. To address this gap, we conducted a pilot EEG experiment testing whether natural soundscapes buffer the neural dysregulation induced by environmental noise. Twelve healthy adults completed a battery of cognitive tasks under three auditory contexts: (1) Noise (urban traffic), (2) Nature (ambient natural sounds), and (3) Control (silence with earplugs). Using spectral, ratio-based, and nonlinear EEG features, we examined how auditory environments modulate cortical efficiency, cognitive load, and affective attention. We hypothesized that (i) noise would elicit signatures of cognitive strain, decreased posterior *α* power, elevated *β* and *θ/β* ratios, and increased entropy, while (ii) nature would preserve balanced oscillatory dynamics indicative of relaxation and attentional stability. This work advances environmental neuroscience by integrating electrophysiological, behavioral, and health metrics to reveal how everyday auditory contexts shape the brain’s capacity for efficient and adaptive regulation.

## 2 Methods

### 2.1 Participants

Twelve healthy adults (4 females, 8 males; ages 19 to 49) were recruited for this pilot study. All participants were office workers or students with normal hearing and no known neurological conditions. The sample size was determined by the practical constraints of this preliminary work, as the goal was to obtain initial effect-size estimates for future, larger studies. Participants provided written informed consent. The study protocol was reviewed and approved by an independent Institutional Review Board (Advarra IRB) and was conducted under the oversight of the Xheladin and Xhufe Morina Foundation (XhMF) and its research arm, the Environmental Neuroscience Research Incubator (ENR Index). All experimental sessions took place in Boston, Massachusetts. A within-subjects experimental design was employed in which each participant completed three environmental conditions: Noise, Nature, and Control. Each condition lasted approximately 20 minutes, for a total session time of about 60 minutes per participant. The order of conditions and tasks was counterbalanced across participants to control for fatigue and practice effects. Testing was conducted individually in a quiet room resembling a typical office environment. For each condition, participants sat at a desk equipped with a monitor, separate speakers, a keyboard, and an air-quality sensor, where they completed the surveys and cognitive tasks (Figure 1). In the Noise condition, participants were exposed to a recorded urban traffic soundscape while performing the tasks. This soundscape included a mix of common urban noises such as distant conversations and traffic hum, averaging approximately 70dB(A) with occasional peaks near 75dB. The recording was designed to simulate a busy yet realistic environment with unpredictable salient events but without any abrupt or startling sounds above 80dB. In the Nature condition, participants performed the same tasks while listening to a continuous natural soundscape consisting of gentle outdoor ambient noises (rustling leaves, birdsong, and flowing water) at the same average intensity level (70dB(A)). The natural recording was selected to be continuous and soothing, lacking sharp acoustic transients. Playback calibration was verified with NIOSH SLM app sound level meter, using an iPhone. Participants were informed that the background sounds were not part of the task and were instructed simply to perform as they normally would. They were not informed of the study’s hypotheses until debriefing. Each condition block (Noise, Nature, or Control) consisted of three segments: (1) a 2-minute baseline period to rest quietly and acclimate to the soundscape, (2) approximately 10 to 12 minutes of cognitive task performance, and (3) a brief post-condition survey assessing subjective experience. Before beginning the first condition, participants also completed a demographic questionnaire and baseline mood and stress measures for reference.

**Figure 1.**
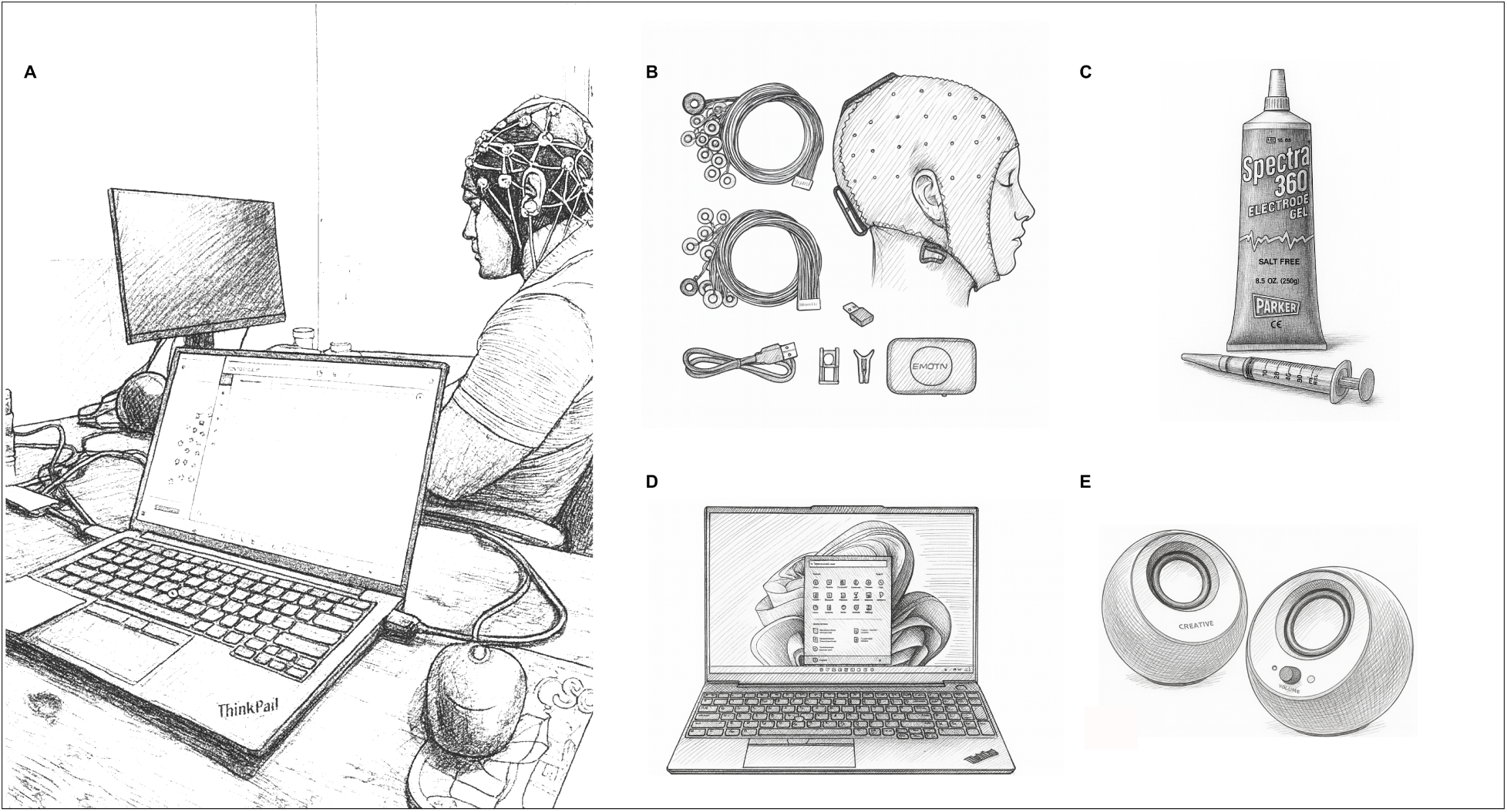
Experimental setup and components of EEG acquisition. (A) Participant seated in a quiet, office-like testing room while completing cognitive tasks on a computer monitor, wearing a 32-channel EEG cap. (B) Components of the EEG acquisition system, including the electrode cap, amplifier, and connection cables. (C) Conductive materials (Spectra 360 electrode gel and syringe applicator) used for electrode preparation. (D) Experimenter’s laptop displaying the real-time EEG acquisition interface used to monitor signal quality and mark task time frames. (E) External stereo speakers delivering the auditory environments (Noise, Nature, Control) at matched sound-pressure levels.

**Figure 2.**
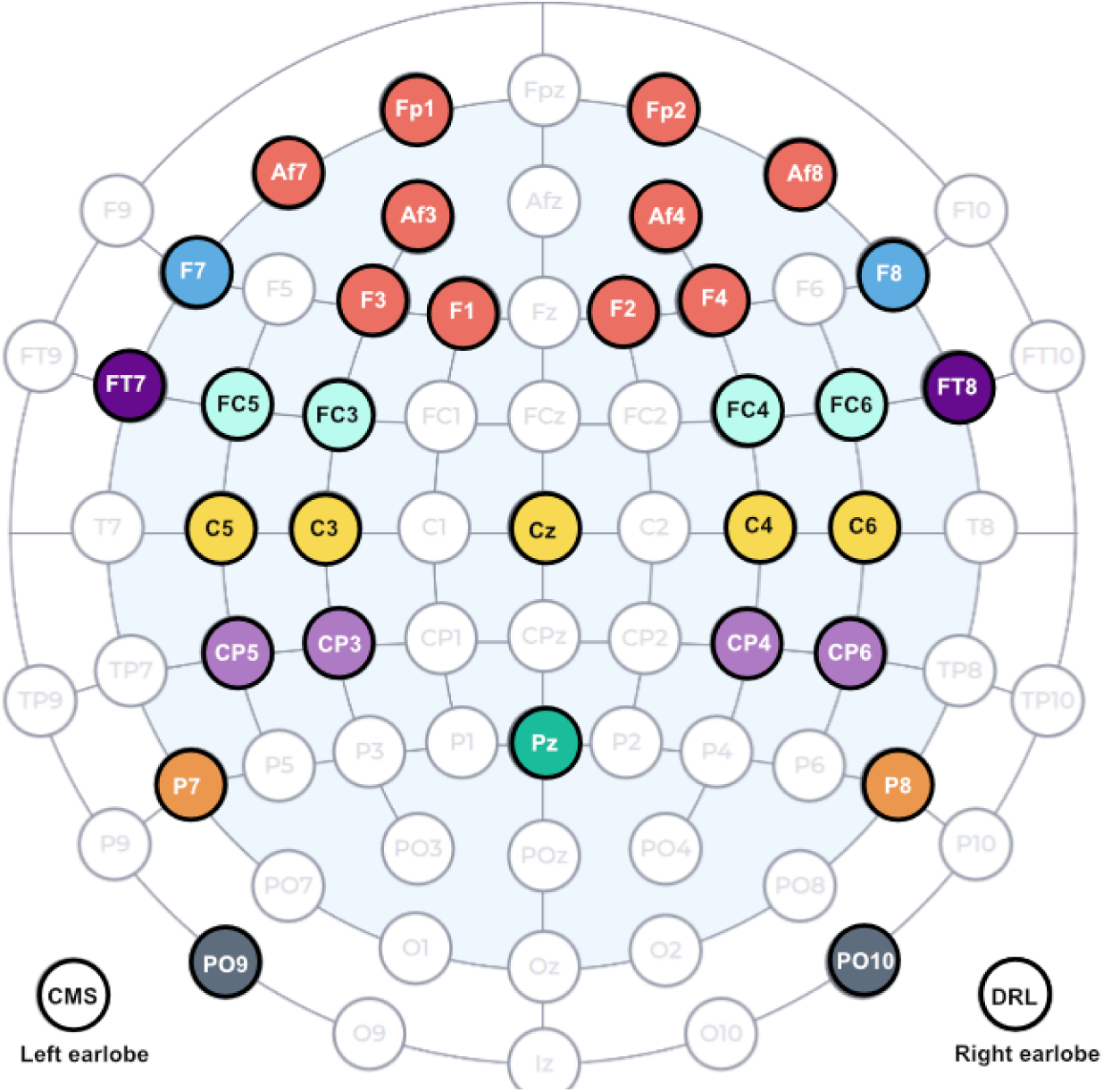
EEG electrode placement and grouping for analysis. Thirty-two electrodes were positioned according to the extended international 10–20 system. The schematic illustrates the spatial distribution of electrodes and their assignment to nine regions of interest (ROIs) based on spatial proximity and functional relevance: Frontal (10 sensors) Fp1, Fp2, F1, F2, F3, F4, Af3, Af4, Af7, Af8; Frontal Lateral (2 sensors) F7, F8; fronto-central (4 sensors) FC3, FC4, FC5, FC6; Central (5 sensors) C3, C4, C5, C6, Cz; centro-parietal (4 sensors) CP3, CP4, CP5, CP6; parietal (1 sensor) Pz; parietal lateral (2 sensors) P7, P8; parieto-occipital (2 sensors) PO9, PO10; Temporal (2 sensors) FT7, FT8. Data from electrodes within each ROI were averaged to reduce dimensionality and to examine regional effects of environmental condition on neural activity.

### 2.2 Cognitive Task Battery

During each auditory condition, participants completed a standardized battery of computerized tasks assessing attention, inhibition, and working memory. All paradigms were adapted from validated versions implemented in PsyToolkit **?**and presented via a custom interface on a computer monitor.

- NBack Task (working memory). Participants performed a 2-back letter task requiring continuous monitoring and updating of stimuli. Uppercase letters (A-Z) appeared sequentially at the center of the screen (stimulus duration = 500ms; inter-stimulus interval = 2500ms). Participants pressed the space bar when the current letter matched the one presented two trials earlier. Accuracy (percentage of targets correctly identified), d-prime (signal-detection sensitivity), criterion (response bias), and reaction time (RT in ms) for correct trials were extracted.
- DotProbe Task (attention bias). Each trial displayed an emotional-neutral word pair followed by a probe (dot) replacing one of the words. Participants indicated the probe position (left or right) as quickly as possible. Reaction times for emotional versus neutral trials were used to compute an attention-bias index = (incongruent RT - congruent RT).
- Visual Search Task (visual attention). Participants searched for an inverted letter T among upright lettr T distractors. Accuracy (percentage correct) and RT were recorded. Search efficiency (RT per set size) and slope parameters were derived as indices of visual attention and resistance to distraction.
- Stroop Task (cognitive control). Color words (e.g., red, green, blue) were presented in congruent or incongruent ink colors. Participants responded to the ink color. Accuracy (percentage correct) and RT (ms) were computed for each trial type, and the Stroop effect = (incongruent RT - congruent RT) quantified interference control.
- GoNoGo Task (response inhibition). Sequential letters appeared on the screen. Participants responded to Go stimuli (80% of trials) and withheld responses to No Go stimuli (20%). Go-trial RTs and commission errors (false alarms) were analyzed as measures of inhibitory control and sustained attention.

For each participant, task performance was summarized separately for the Control, Nature, and Noise conditions. Within-subject differences across conditions were calculated for all dependent variables. Primary behavioral outcomes included cognitive control (Stroop effect), attention bias (DotProbe index), inhibitory control (GoNoGo performance), working memory (NBack accuracy and d-prime), and visual attention (Visual Search efficiency). EEG-derived measures (described below) were analyzed in parallel to identify neural correlates of environmental modulation.

### 2.3 EEG Recordings

#### 2.3.1 EEG Acquisition and Preprocessing

Continuous EEG was recorded throughout each auditory condition using a 32-channel wireless headset (EMOTIV Flex Gel; sampling rate = 256Hz). Electrodes were positioned according to the extended international 10-20 system (Figure2), providing coverage across nine regions of interest (ROIs). Signals were referenced to the common average, and electrode impedances were maintained below 10kΩ. Data were band-pass filtered between 0.5 and 100Hz using a zero-phase finite impulse response (FIR) filter and subjected to automated quality-control checks. Channels with a standard deviation below 0.5*µ*V (flat) or above 150*µ*V (noisy) were flagged and excluded. Ocular and muscle artifacts were removed using independent component analysis (ICA) after visual inspection of components. The cleaned EEG signals were segmented into baseline (rest) and task periods using 2-second epochs with 50% temporal overlap for each auditory condition (Noise, Nature, and Control). Epochs containing residual artifacts exceeding 100*µ*V were rejected prior to spectral decomposition.

#### 2.3.2 Spectral Analysis and Feature Extraction

Power spectral density (PSD) was computed for each epoch using Welch’s method (2-second windows, 50% overlap). Mean spectral power was calculated within canonical frequency bands: *δ* (0.5-4Hz), *θ* (4-7Hz), *α* (8-12Hz), *β* (12-30Hz), and *γ* (30-100Hz). Power values were log_10_-transformed, z-score normalized within participants, and averaged across electrodes within nine regions of interest (ROIs): central, centro-parietal, frontal, fronto-central, frontal-lateral, parietal, parietal-lateral, parieto-occipital, and temporal. From these spectra, several indices of cognitive and affective processing were derived:

- Frontal Alpha Asymmetry (FAA). FAA was computed by averaging alpha power across left and right frontal hemispheres for each condition and task, taking the log_10_-transformed difference (Right - Left), standardizing within participants (z-score), and then computing the baseline-corrected difference (Task - Baseline). Positive FAA values indicate greater relative right-frontal activation (approach motivation or positive affect), whereas negative values suggest withdrawal or stress-related activation.

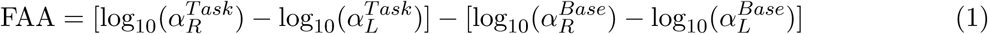
- Theta/Alpha Ratio. The log-transformed ratio log_10_(*θ*) − log_10_(*α*) was computed for each channel, averaged within ROIs, standardized within participants, and baseline-corrected (Task - Baseline). Higher values reflect increased cognitive load and mental effort relative to baseline.

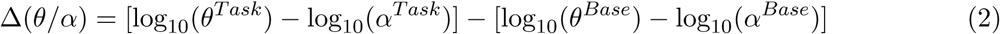
- Theta/Beta Ratio. The log-transformed ratio log_10_(*θ*) − log_10_(*β*) was computed for each channel, averaged within ROIs, standardized within participants, and baseline-corrected (Task - Baseline). Elevated theta/beta ratios are associated with reduced attentional stability and cognitive dysregulation.

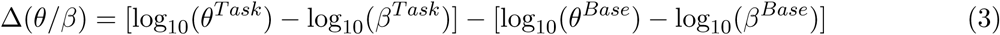
- Engagement Index (EI). EI was computed as the log_10_-transformed ratio of (*β* + *γ*) power to (*α* + *θ*) power for each channel, averaged within ROIs, standardized within participants, and baseline-corrected (Task - Baseline). Higher values indicate increased cortical activation and task engagement relative to baseline.

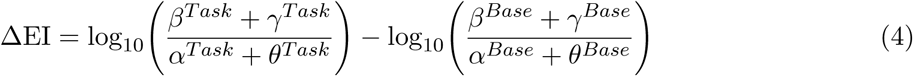

To further characterize neural dynamics beyond spectral power, additional time-domain and nonlinear features were extracted from each 2-second epoch (Lim et al., 2022), including skewness, Hjorth mobility, Hurst exponent, detrended fluctuation analysis (DFA), and sample entropy.

#### 2.3.3 Statistical Analysis

EEG band power and derived indices were analyzed using linear mixed-effects models (LMMs) implemented in Python (Statsmodels). Condition was specified as a fixed effect, and participant as a random intercept to account for repeated measures. Condition order and task order were fully counterbalanced across participants. To control for Type I error across nine ROIs and five frequency bands (45 tests per metric), *p*-values were adjusted using the Benjamini-Hochberg false discovery rate (FDR) procedure. Statistical significance was defined at *p <* 0.05 and *p <* 0.01 for highly significant effects. Effect sizes were quantified using *t*-statistics from the LMMs. Internal reliability of spectral measures was evaluated using intra-class correlations (ICC[3,1]) and split-half reliability (Spearman-Brown corrected) computed for each condition. Mean ICCs and split-half coefficients are reported in the Results section to support measurement stability.

### 2.4 Environmental and Neurological Risk (ENR) Index

To assess individual differences in baseline health status that might moderate neural responses to environmental context, we computed an Environmental and Neurological Risk (ENR) Index for each participant. The ENR Index is a composite score derived from self-reported health and lifestyle factors collected via questionnaire at the start of the experimental session. The index integrates 11 health domains: environmental exposure (based on zip code), blood pressure, physical activity, diet quality, sleep quality, smoking status, social engagement, perceived stress levels, indoor ventilation quality, medication use, substance use, and pre-existing health conditions. Each domain was scored on a standardized scale (typically 0-3, where higher values indicate better health), and domain scores were summed to create a total ENR Index score. Higher ENR Index scores reflect better overall health and lower environmental risk. Scores were categorized into three risk levels: Good health (ENR ≥ 14), Moderate health (7.5 ≤ ENR *<* 14), and Low health (ENR *<* 7.5). The ENR Index was used as a continuous moderator variable in correlation analyses examining whether baseline health status predicted individual differences in neural responses to Nature and Noise conditions. This tool was inspired by the BrainCare Score, used to measure dementia risk (Singh et al., 2023). (See Supplementary Figure 2.)

## 3 Results

### 3.1 Neural Complexity and Signal Dynamics Across Environments

To assess how auditory environments modulated intrinsic neural dynamics, we first examined nonlinear EEG features that reflect temporal complexity and statistical regularity of the signal (Figure 3 and 4). Across most metrics, neural activity exhibited strong stability across environmental conditions. Detrended Fluctuation Analysis (DFA), sample entropy, and Hurst exponent values remained consistent across Control, Nature, and Noise, with regional averages ranging from 0.7 to 1.2. Fronto-central electrodes showed the highest values (∼ 0.8-1.0), whereas parieto-occipital and parietal-lateral regions consistently displayed slightly lower values. Skewness demonstrated region-specific asymmetries: frontal-lateral and temporal regions exhibited negative skewness (−0.02 to −0.05), while parietal and parieto-occipital regions showed positive skewness (≈ 0.02 to +0.06) in the Nature condition. In the Control condition, frontal-lateral, parietal-lateral, and temporal regions again showed negative skewness, a pattern largely replicated under Noise. Hjorth mobility revealed a modest environmental modulation, with higher mobility observed in the central region during the Nature condition (0.37) compared with Control and Noise (∼ 0.30-0.35). Other regions displayed comparable values across conditions. Overall, nonlinear EEG measures indicated that environmental effects were subtle and spatially restricted. Temporal dynamics of neural activity remained robust to environmental change, with the Nature condition producing the most distinguishable, yet localized, alterations, and Noise exerting minimal disruption on intrinsic temporal organization.

**Figure 3.**
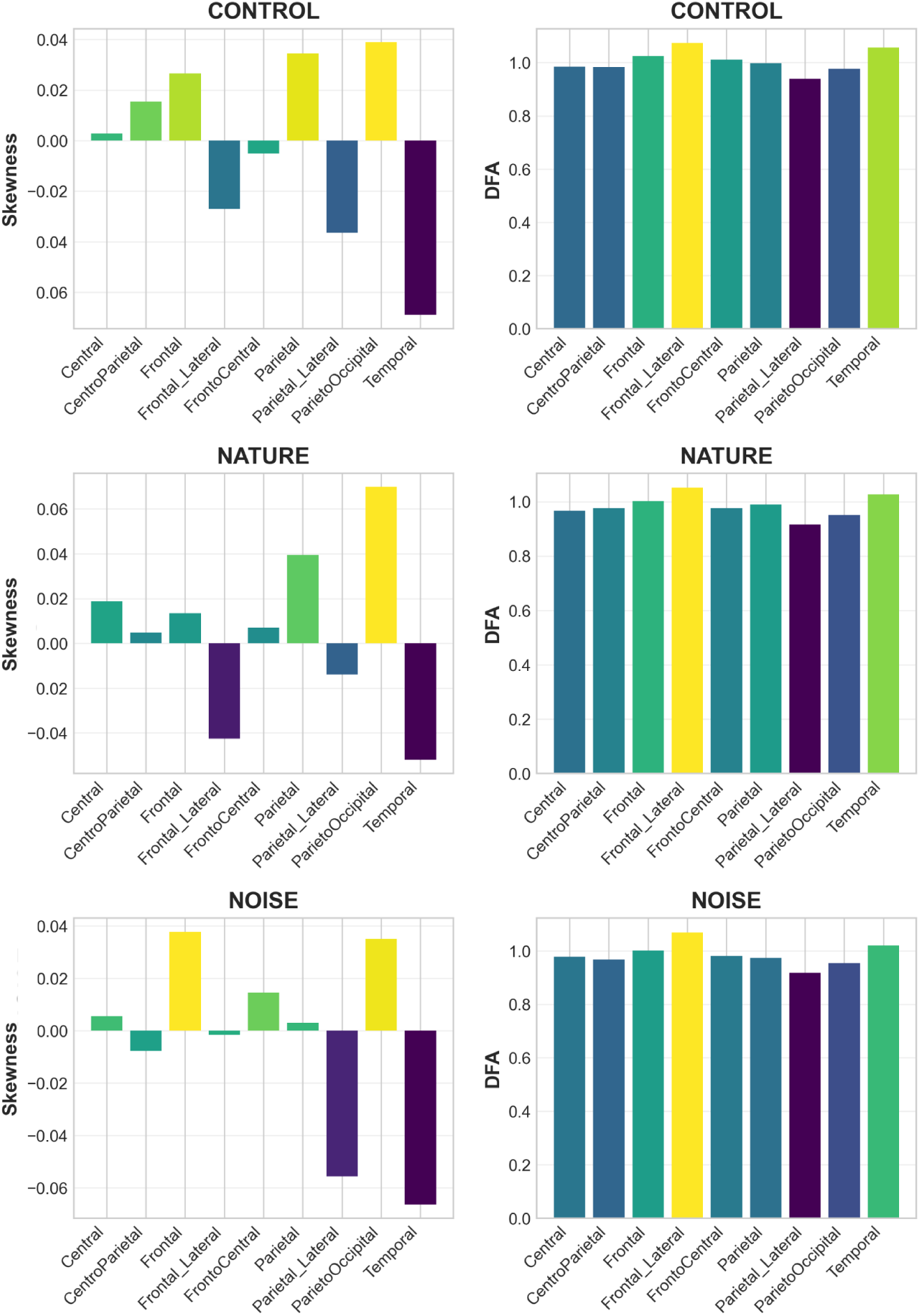
Time-domain and nonlinear dynamics. Mean values of Skewness (left column) and Detrended Fluctuation Analysis (DFA) (right column) are shown for each region of interest (ROI) across the three auditory conditions: Control (top row), Nature (middle row), and Noise (bottom row).

**Figure 4.**
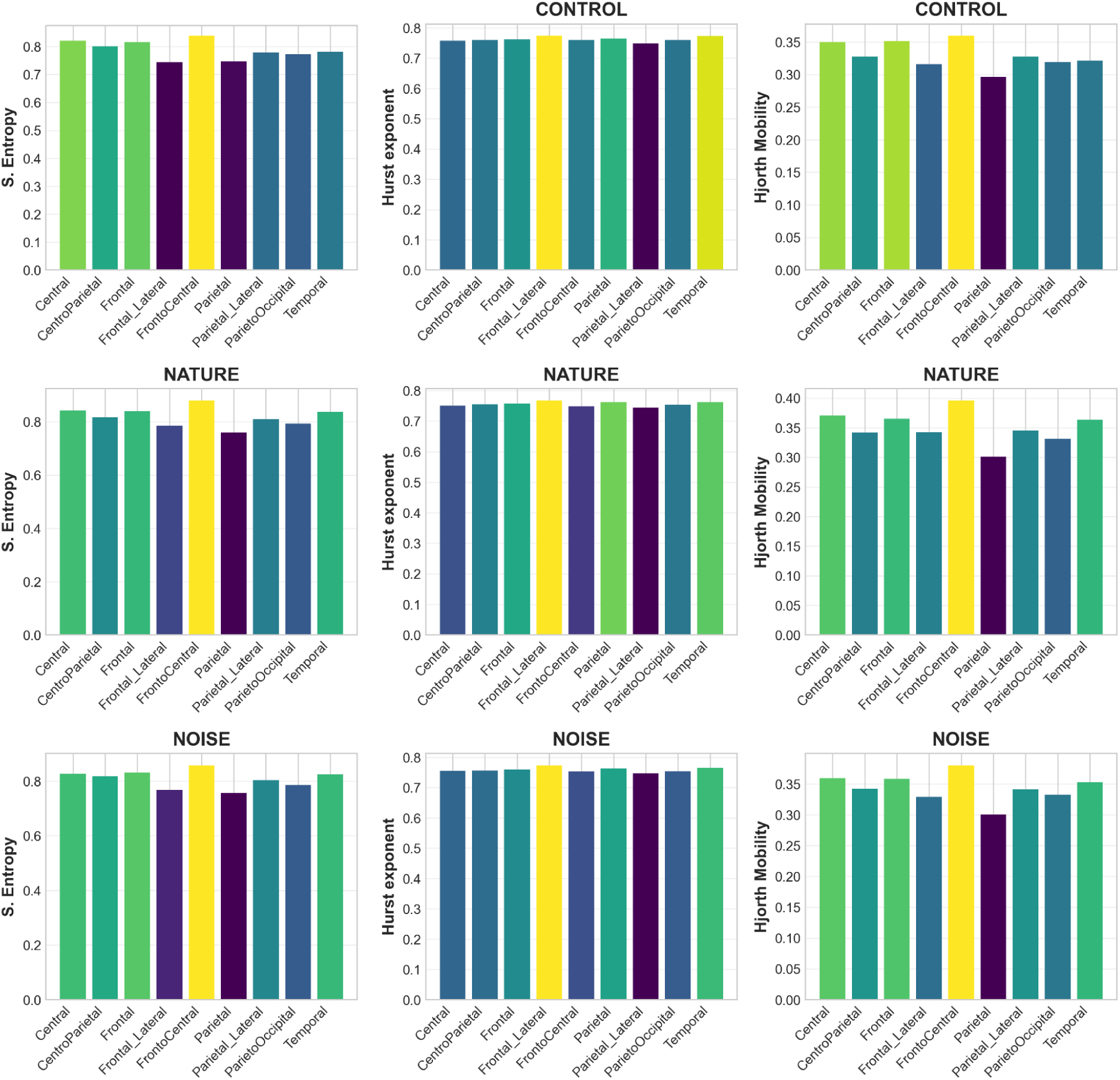
Complexity features and temporal organization. Mean values of Spectral Entropy (left column), Hurst Exponent (middle column), and Hjorth Mobility (right column) are shown for each region of interest (ROI) across the three auditory conditions: Control (top row), Nature (middle row), and Noise (bottom row).

### 3.2 Neural Activity in Noise versus Nature

#### 3.2.1 Spectral Power and Cognitive Indices

Spectral analyses with false-discovery-rate (FDR) correction revealed distinct neural signatures differentiating the Noise and Nature conditions across multiple frequency bands and cortical regions. Linear mixed-effects modeling identified 17 significant condition effects (FDR-corrected *p <* 0.05) spanning spectral and ratio-based indices (Figure5). As predicted, exposure to Noise elicited patterns consistent with higher cognitive strain and arousal, whereas Nature soundscapes were associated with more regulated cortical activity. The most robust environmental effects emerged in the *δ* band, which exhibited widespread modulation across five ROIs (centro-parietal, parietal, parietal-lateral, parieto-occipital, and fronto-central), peaking in parietal-lateral regions (− log_10_ *p* ≈ 2.1). This finding suggests that slow-frequency oscillations, typically linked to large-scale cortical coordination, are sensitive to environmental context even during active cognitive engagement. Cognitive ratio indices also showed strong posterior effects. Both the *θ/β* and *θ/α* ratios increased significantly in the parieto-occipital ROI (− log_10_ *p* ≈ 2.1 for each), indicating heightened cognitive-load markers within posterior attention networks. The *α/β* ratio revealed significant modulation in parietal (− log_10_ *p* ≈ 2.0) and fronto-central (− log_10_ *p* ≈ 1.6) regions, reflecting shifts in the balance between relaxed (alpha) and alert (beta) states. At higher frequencies, *β* power was significantly modulated in parietal-lateral and parieto-occipital regions (− log_10_ *p* ≈ 1.6), while *γ* power showed comparable effects in parietal-lateral and fronto-central areas (− log_10_ *p* ≈ 1.6). These findings point to environment-dependent adjustments in cortical synchronization and alertness. Finally, the Engagement Index, representing the ratio of high- to low-frequency activity, was significantly influenced by condition in the parieto-occipital and fronto-central regions (− log_10_ *p* ≈ 1.6). This pattern indicates that environmental context modulates large-scale engagement across posterior attention and frontal executive networks. Together, these results demonstrate a posterior-dominant pattern of environmental sensitivity, with the strongest and most consistent effects localized to parieto-occipital and parietal-lateral regions. These areas, implicated in sensory integration and attentional control, appear more responsive to auditory environmental context than primary motor or purely frontal executive regions.

#### 3.2.2 Direct Spectral Power Analysis

Direct comparisons of spectral power across auditory environments revealed distinct, condition-specific modulation patterns (Figure 6). The upper panels depict contrasts between each experimental condition and Control, while the lower panels compare Nature and Noise directly.

- *Nature vs. Control* Exposure to Nature soundscapes produced selective increases in spectral power relative to the Control condition. Delta-band activity was significantly elevated across four posterior regions, centro-parietal (*p <* 0.05), parietal (*p <* 0.05), parietal-lateral (*p <* 0.01), and parieto-occipital (*p <* 0.05), with the strongest effect in parietal-lateral cortex. Beta power increased in fronto-central (*p <* 0.05), centro-parietal (*p <* 0.05), and parietal-lateral (*p <* 0.05) regions. Gamma-band activity also rose significantly in four regions, frontal-lateral (*p <* 0.05), fronto-central (*p <* 0.05), parietal-lateral (*p <* 0.01), and parieto-occipital (*p <* 0.05), again peaking at parietal-lateral sites (*p <* 0.01). No significant differences were observed for alpha or theta bands, indicating that Nature exposure selectively enhanced low-(delta) and high-frequency (beta, gamma) oscillations while leaving mid-frequency rhythms largely stable.
- *Noise vs. Control.* Noise exposure produced a broader and more spatially extensive pattern of spectral modulation. Delta power increased significantly across seven cortical regions, frontal (*p <* 0.05), fronto-central (*p <* 0.05), central (*p <* 0.05), centro-parietal (*p <* 0.05), parietal (*p <* 0.05), parietal-lateral (*p <* 0.01), and parieto-occipital (*p <* 0.01∗), representing the most widespread delta enhancement observed. The strongest effects again localized to posterior cortices (parietal-lateral and parieto-occipital, *p <* 0.01). Theta power was significantly elevated in the parietal-lateral region (*p <* 0.01), reflecting increased low-frequency activity in attention-related areas. Beta power showed moderate increases in frontal and parietal-lateral regions (*p <* 0.05), whereas gamma power remained unchanged relative to Control, suggesting that Noise primarily accentuates slower oscillatory dynamics.
- *Noise vs. Nature.* Direct contrasts between Noise and Nature conditions (Figure 6, lower panels) revealed that Noise induced significantly higher delta power in four cortical regions, central (*p <* 0.05), centro-parietal (*p <* 0.05), parietal-lateral (*p <* 0.01), and parieto-occipital (*p <* 0.01). No significant differences were detected for theta, alpha, beta, or gamma bands, indicating that delta oscillations represent the principal spectral domain differentiating the two environmental contexts. Collectively, these findings suggest that Noise exposure enhances slow-wave (delta) synchronization, consistent with increased cognitive effort or cortical strain, whereas Nature exposure promotes a balanced oscillatory profile, with selective reinforcement of delta, beta, and gamma activity supporting relaxed yet engaged neural states.

**Figure 5.**
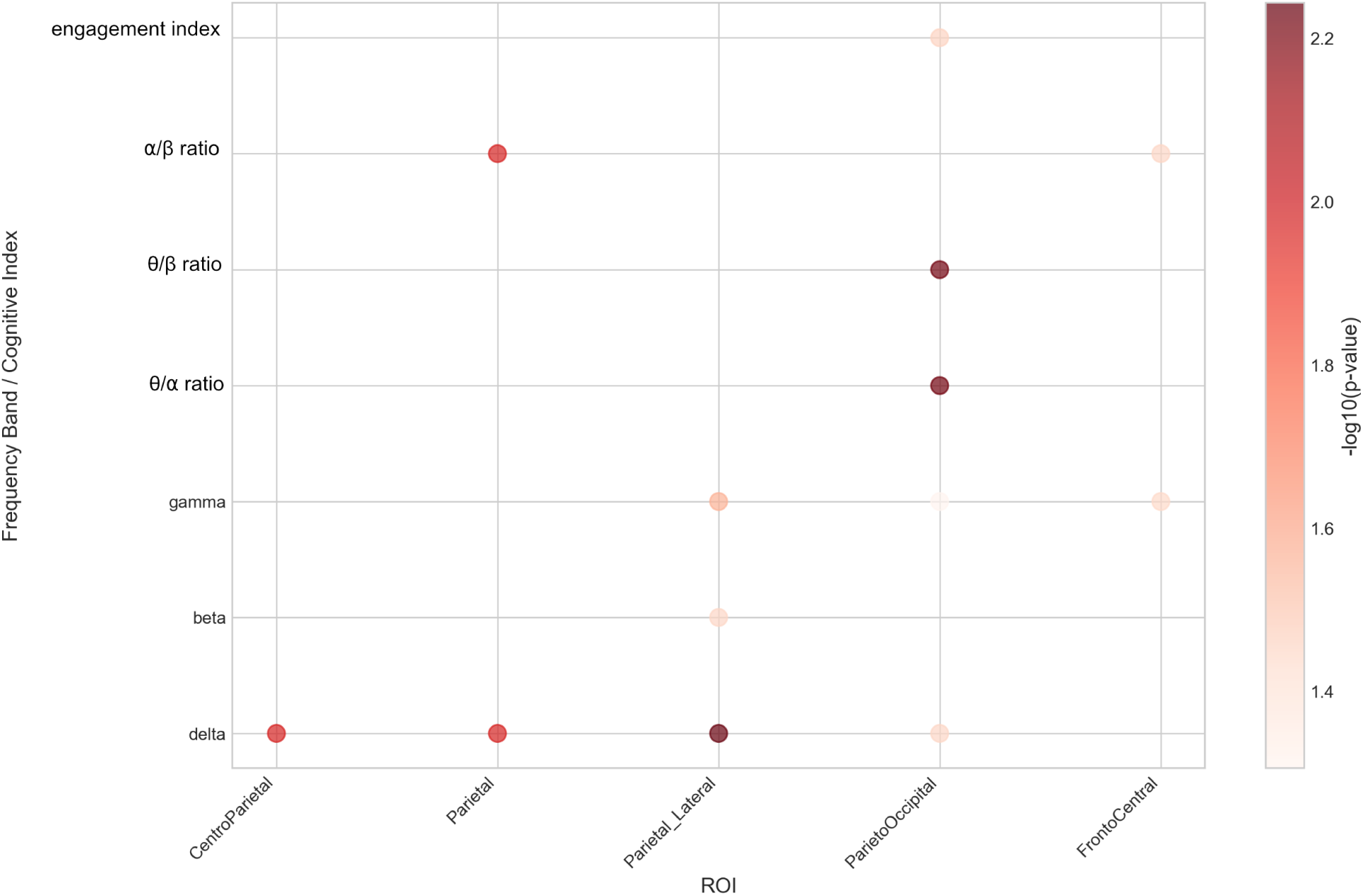
Significant Mixed-Effects Results. This heatmap shows −*log*_10_ transformed p-values from mixed-effects models testing condition effects (Noise, Nature, Control) across frequency bands and regions of interest (ROIs). Each cell plots a circular marker colored by −*log*_10_(p-value), where darker red indicates greater significance. Y-axis: frequency bands and cognitive ratio indices. X-axis: ROIs, eight brain regions.

**Figure 6.**
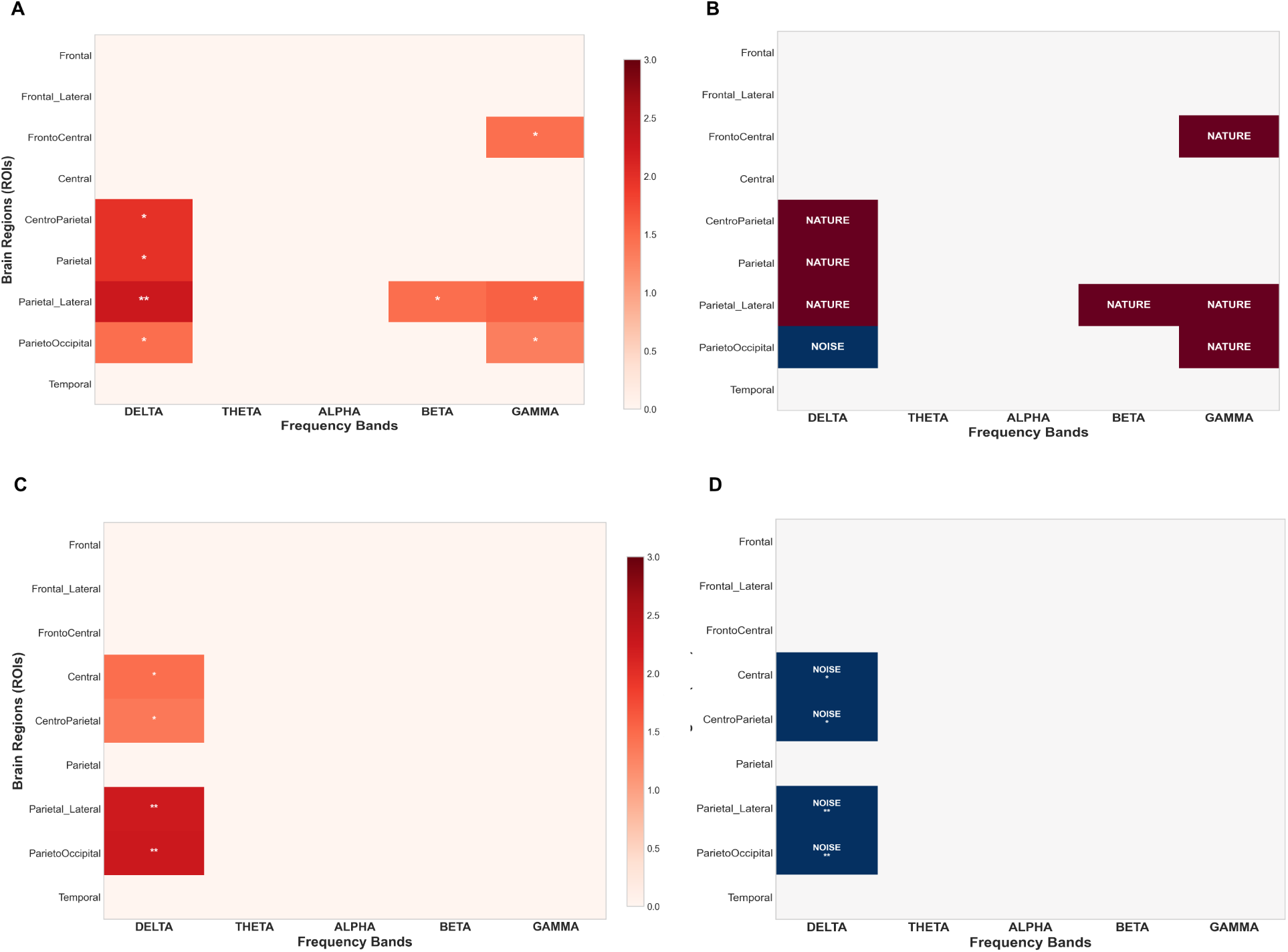
Condition Comparisons: Spectral Band and Region Effects. (A) Group-level results comparing Nature vs. Control and Noise vs. Control conditions across canonical EEG frequency bands and regions of interest (ROIs). Map from mixed-effects analyses, with asterisks marking statistical significance: ∗*p <* 0.05, ∗ ∗ *p <* 0.01, ∗ ∗ ∗*p <* 0.001. Color intensity indicates the strength of the effect. (B) Condition-specific directionality maps, where red = increased spectral power and blue = decreased power relative to the Control condition. (C) Direct comparison of Nature vs. Noise conditions across EEG frequency bands and regions of interest (ROIs). Map from mixed-effects analyses, with asterisks marking statistical significance: ∗*p <* 0.05, ∗ ∗ *p <* 0.01, ∗ ∗ ∗*p <* 0.001. Color intensity indicates the strength of the effect. (D) Condition directionality plot, where red = higher under Nature and blue = higher under Noise.

#### 3.2.3 EEG Band-Ratio Analyses

EEG band-ratio analyses (Figure 7) revealed condition-dependent reorganization of large-scale cortical dynamics across four key cognitive indices. Relative to the Control condition, both Nature and Noise exposures produced selective bilateral effects (“B” in Figure 7), reflecting distinct patterns of neural modulation.

- *Alpha/Beta Ratio (Relaxation vs. Alertness).* Nature exposure significantly increased the *α/β* ratio bilaterally in fronto-central (*p <* 0.05) and parietal (*p <* 0.05) regions (Figure 7D). Elevated *α/β* ratios indicate a shift toward a more relaxed cortical state while preserving executive and attentional regulation. The bilateral pattern suggests coordinated engagement of left- and right-hemisphere networks involved in cognitive control and sustained attention. In contrast, Noise exposure produced no significant *α/β* changes relative to Control, indicating that noise environments do not elicit the same relaxed-yet-alert neural profile observed with Nature soundscapes.
- *Theta/Alpha Ratio (Cognitive Load).* Noise exposure selectively increased *θ/α* ratios bilaterally in parieto-occipital regions (*p <* 0.01; Figure 7A), consistent with elevated cognitive load and more effortful processing. This posterior network, implicated in visuospatial attention, appears particularly sensitive to noise-induced cognitive strain. Nature exposure, by comparison, did not significantly alter *θ/α* ratios, suggesting preserved processing efficiency under restorative auditory conditions.
- *Theta/Beta Ratio (Attentional Regulation).* Noise exposure also significantly reduced *θ/β* ratios bilaterally in parieto-occipital regions (*p <* 0.01; Figure 7B). Such reductions may reflect compensatory adjustments or altered attentional regulation in response to environmental challenge. The co-occurrence of increased *θ/α* and decreased *θ/β* ratios in the same regions points to complex modulation of slow-wave activity, consistent with heightened attentional effort under Noise. Nature exposure again showed no significant *θ/β* changes relative to Control.
- *Engagement Index (β/α Ratio).* Nature exposure produced a bilateral increase in the Engagement Index in parieto-occipital regions (*p <* 0.05; Figure 7C), indicating enhanced cortical activation and task engagement relative to Control. The bilateral enhancement mirrors the *α/β* ratio pattern, together suggesting that Nature soundscapes promote an “engaged relaxation” state, heightened alertness with reduced strain. Noise exposure did not significantly affect the Engagement Index.

**Figure 7.**
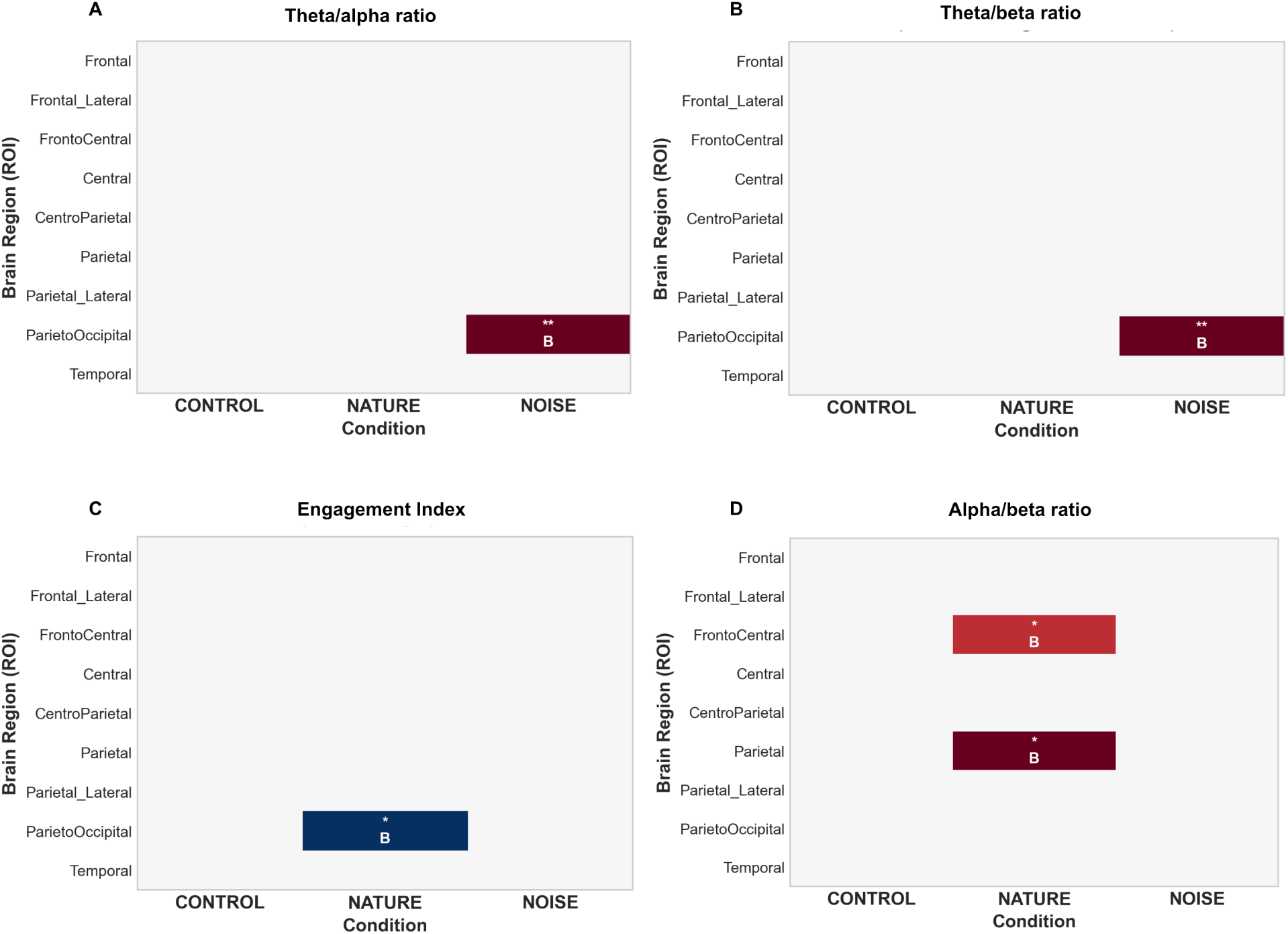
Regional Effects of Environmental Context on Baseline-Corrected Cognitive Indices. Baseline-corrected EEG-derived cognitive indices across nine brain regions (ROIs) for Control, Nature, and Noise conditions. Each heatmap shows the magnitude and direction of effects relative to baseline (color scale: red = increase, blue = decrease relative to baseline). Asterisks denote statistical significance: ∗*p <* 0.05, ∗ ∗ *p <* 0.01, ∗ ∗ ∗*p <* 0.001. The letter “B” indicates bilateral effects. (A) *θ/α* ratio, attention and cognitive load. (B) *θ/β* ratio, ADHD and cognitive control. (C) Engagement Index (EI), cognitive engagement. (D) *α/β* ratio, relaxation vs. alertness.

Collectively, these results reveal distinct environmental profiles. Nature exposure is characterized by bilateral increases in *α/β* ratio (fronto-central and parietal) and Engagement Index (parieto-occipital), consistent with relaxed yet engaged neural states. Noise exposure, in contrast, yields bilateral increases in *θ/α* ratio and reductions in *θ/β* ratio (parieto-occipital), indicating elevated cognitive load and disrupted attentional regulation. The predominance of bilateral effects within posterior attention networks underscores the sensitivity of parieto-occipital regions to auditory environmental context and suggests that environmental modulation of cortical dynamics operates through coordinated, rather than lateralized, network adjustments. Altogether, figures 5, 6 and 7 provide complementary perspectives on how auditory environments shape cortical dynamics. Figure 5 summarizes all significant mixed-effects findings across frequency bands and cognitive indices, Figure 6 presents direct spectral-power contrasts (*δ*, *θ*, *α*, *β*, *γ*) for each condition relative to Control and between Noise and Nature, and Figure 7 focuses on composite cognitive ratios (*α/β*, *θ/α*, *θ/β*, Engagement Index) that integrate multiple frequency bands to reflect functional neural states. Although effects were systematic and spatially distributed, they did not form contiguous clusters, suggesting region-specific rather than global synchronization changes. Taken together, these findings show that Nature soundscapes promote a state of relaxed alertness, characterized by balanced activation across hemispheres and frequency bands, whereas Noise induces a state of cortical strain, marked by elevated slow-wave activity and disrupted attentional control. The convergence across Figures 5-7 highlights that environmental modulation of brain activity is most pronounced in posterior attention networks and in the coupling between low- and high-frequency oscillations.

#### 3.2.4 Task-Specific EEG Effects

Task-specific analyses of baseline-corrected spectral values (Figure 8) showed that environmental modulation of neural activity varied across cognitive tasks. Baseline correction isolated task-evoked changes from resting-state activity, enabling comparison of nine spectral features across the five tasks.

**Figure 8.**
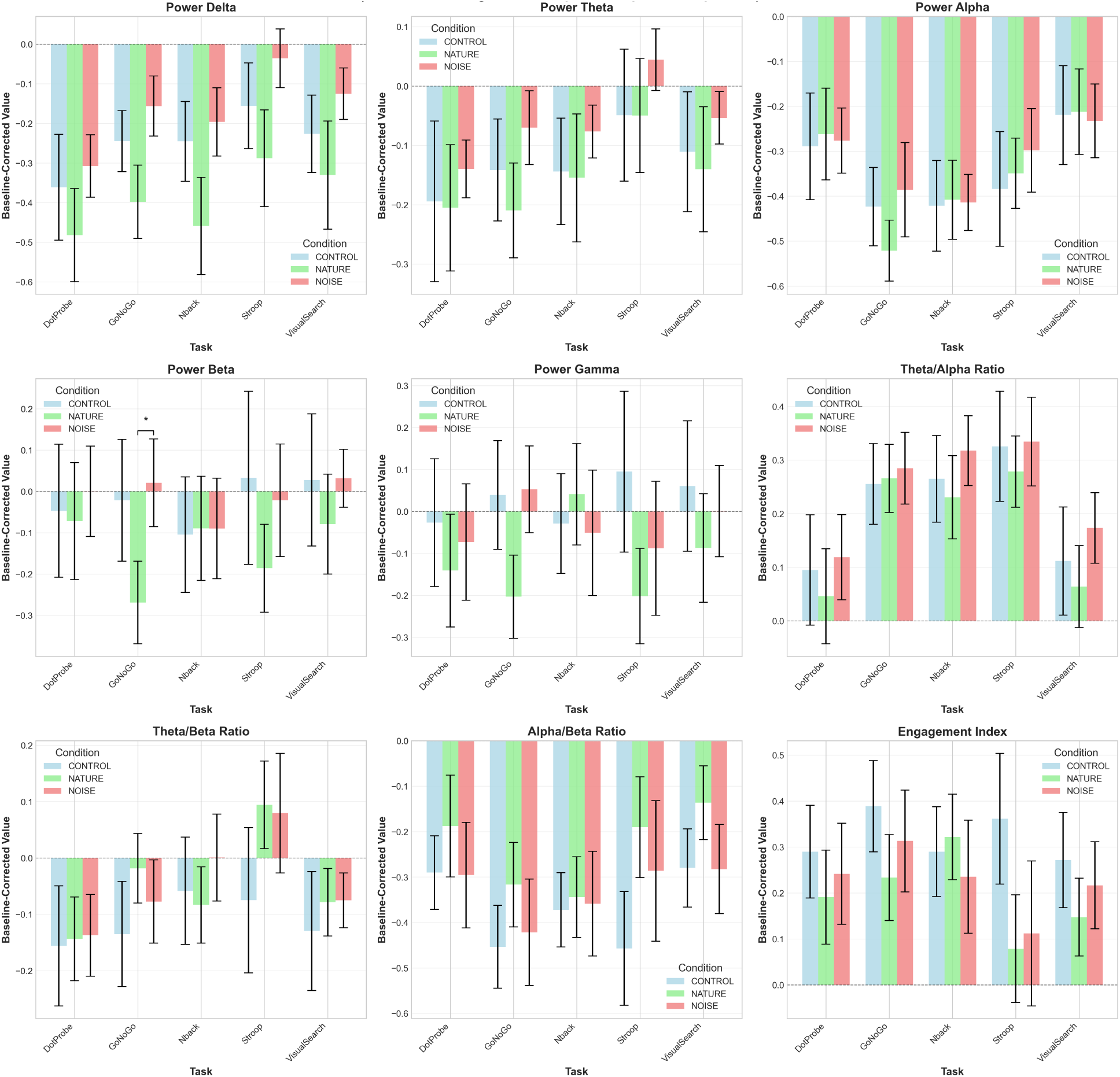
Task-Specific Effects by Condition. Mean baseline-corrected values (task - baseline) for nine EEG features and cognitive ratios across five tasks (DotProbe, GoNoGo, NBack, Stroop, VisualSearch) under three conditions: Control (light blue), Nature (green), Noise (red). Features include spectral power (Delta, Theta, Alpha, Beta, Gamma) and cognitive ratios (Theta/Alpha, Theta/Beta, Alpha/Beta, Engagement Index). Error bars = SEM. Horizontal dashed line = no change from baseline. Asterisks (*) = FDR-corrected *p <* 0.05.

Mixed-effects analyses with FDR correction revealed distinct condition- and task-dependent patterns. The *α/β* ratio exhibited consistently negative baseline-corrected values across all tasks, indicating reductions from baseline during cognitive engagement. However, Nature exposure maintained higher (less negative) *α/β* ratios than Noise in every task, suggesting that Nature preserved a more balanced relaxed-alert state relative to baseline. Beta power showed the most pronounced task-specific differences. During the GoNoGo task, Nature exposure produced substantial beta decreases from baseline, whereas Noise exposure produced modest increases, yielding a significant condition effect (p < 0.05, FDR-corrected). This divergence implies that inhibitory-control demands are met with reduced cortical activation under Nature (reflecting greater efficiency), but with increased beta activity under Noise (indicating heightened effort or strain). Across other tasks, beta changes were smaller and did not differ significantly between conditions. No other spectral features survived FDR correction at the task-specific level, suggesting that environmental influences are concentrated within particular frequency bands.

#### 3.2.5 Cluster-Based Permutation Analysis and Summary

Lastly, cluster-based permutation testing revealed no spatially contiguous clusters surviving FDR correction, indicating that although effects were systematic and distributed across cortical regions, they did not form coherent spatial clusters. Taken together, these findings demonstrate that Nature and Noise exert distinct, large-scale influences on cortical dynamics. Nature exposure was associated with reduced cognitive load (lower *θ/α* in parietal regions), enhanced executive control (lower *θ/β* in fronto-central and parieto-occipital regions), greater relaxation (higher *α/β* across multiple regions), and increased engagement (higher Engagement Index across widespread areas). This constellation of effects suggests more efficient and optimally balanced cortical resource allocation. In contrast, Noise exposure elicited increased cognitive strain (higher *θ/α* in parietal-lateral and parieto-occipital regions), elevated high-frequency power (*δ*, *β*, *γ*), and heightened vigilance without corresponding relaxation benefits, reflecting effortful processing and sustained cortical tension.

### 3.3 Reliability of Baseline Neural Measures

We assessed the internal consistency of EEG spectral power within each auditory baseline using ICC(3,1) computed across non-overlapping 4 s epochs (Figure 9). Reliability differed systematically by environmental context. The Control condition yielded moderate reliability for most bands (*α*-*γ* = 0.66-0.68; *δ*, *θ* = 0.49-0.55). The Noise condition showed the highest stability, reaching 0.73-0.75 for *β*-*γ* (moderate-good range). In contrast, the Nature baseline exhibited markedly lower ICCs (0.28-0.49 across bands), indicating greater temporal variability in neural dynamics. Averaged across all conditions, ICCs ranged from 0.53-0.61, consistent with moderate reliability typical for short resting EEG baselines. This pattern suggests that baseline brain activity is most temporally stable in structured or high-load auditory environments (Control, Noise) and more variable under natural soundscapes. Rather than reflecting noise or poor signal quality, the reduced ICCs in the Nature condition likely index greater spontaneous fluctuation or neural flexibility during restorative states. Importantly, the ratio-based and baseline-corrected indices used in our main analyses (*θ/α*, *θ/β*, *α/β*, Engagement Index) demonstrated higher split-half reliability (median Spearman-Brown corrected *r* = 0.68-0.77), confirming sufficient measurement stability for condition contrasts.

**Figure 9.**
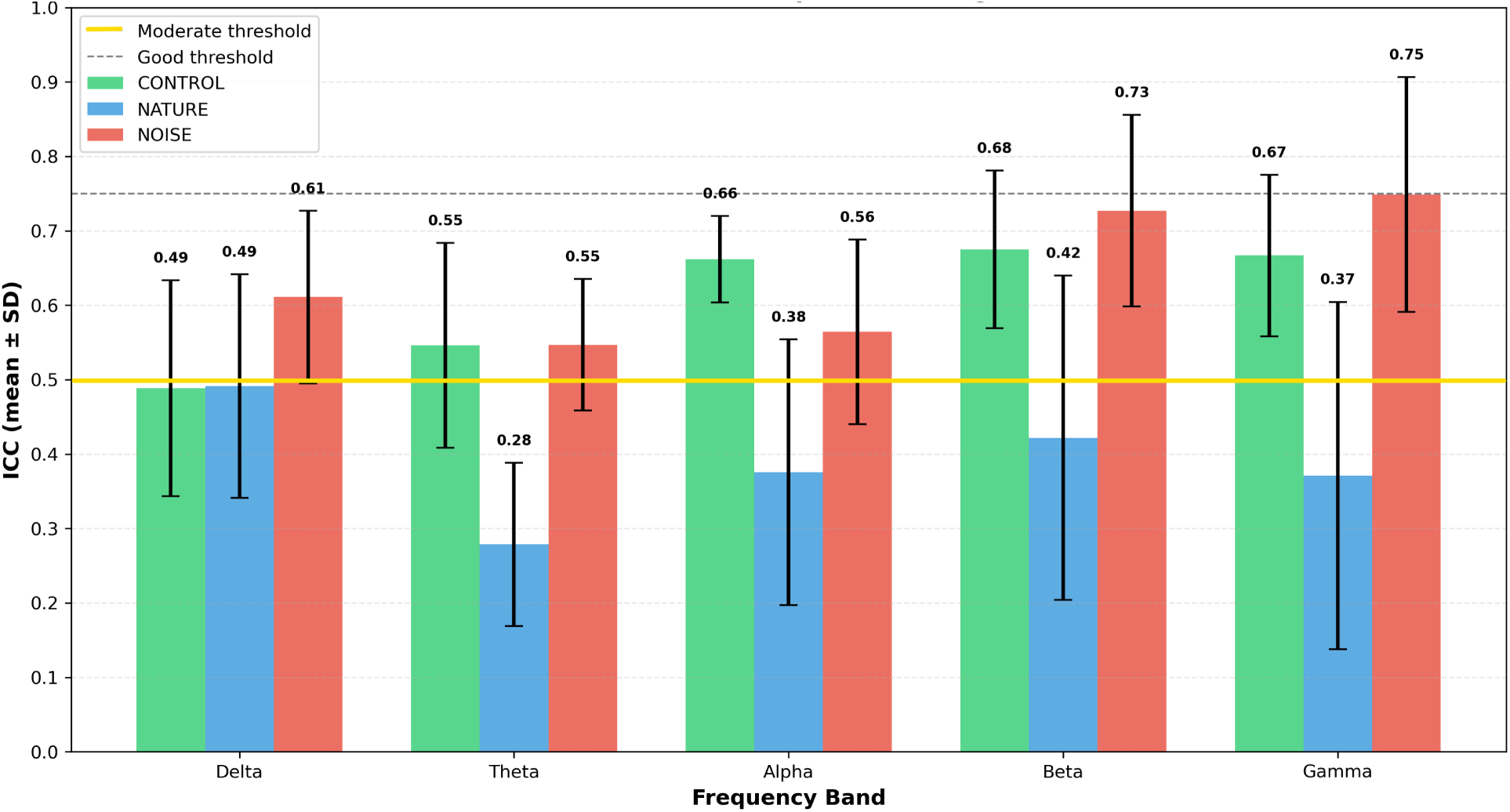
Internal Reliability of Baseline EEG Bands by Condition. Values reflect mean SD across ROIs. As ICC reflects both signal stability and between-subject variance, the reduced reliability in the Nature condition likely represents increased state variability rather than poor signal quality. Importantly, the ratio-based and baseline-corrected indices used for main analyses showed higher split-half reliability (median Spearman-Brown = 0.68-0.77), confirming adequate measurement stability for condition contrasts.

### 3.4 Behavioral Performance

Behavioral outcomes revealed subtle yet meaningful effects of environmental sound on cognitive efficiency and emotional attention. Across all tasks, accuracy remained high (89.9-99.5%), indicating robust overall performance. However, task-specific analyses identified domains most sensitive to environmental manipulation. Performance on the Nback task showed the clearest environmental effects. Mean accuracy improved under Nature (94.4 %) relative to Control (89.9%) and Noise (92.4%), representing a 4.5 % gain consistent with a medium effect size. Noise modestly reduced accuracy relative to Nature, aligning with EEG evidence of elevated frontal *θ/β* ratios and reduced *α* regulation. Stroop and GoNoGo tasks exhibited near-ceiling accuracy (96.0-99.0 %) across all conditions, indicating strong inhibitory control and limited susceptibility to environmental interference. The Visual-Search task yielded the highest accuracy overall (98.9-99.5 %), suggesting that low-demand, perceptually guided tasks are largely insensitive to auditory context. (Figure 9). Then, to evaluate potential effects on affective attention, we examined DotProbe attention-bias indices (Figure 10). Median bias scores were near zero in Control (−2*ms*) and slightly negative in Nature (−8*ms*), indicating balanced or mildly avoidant responses to emotional stimuli. Under Noise, however, the median shifted to +19*ms*, reflecting faster orienting toward emotionally salient or threatening cues. The interquartile range for Noise (+5*to* + 28*ms*) was entirely positive, indicating a consistent attentional bias toward emotional stimuli. Individual trajectories revealed greater variability under Noise, suggesting reduced attentional stability in unpredictable auditory contexts. Overall, behavioral accuracy remained stable across most tasks. Combined with EEG findings, these results indicate a neural-behavioral dissociation: participants maintained high task performance even as cortical activity patterns diverged across environments. This suggests that Noise required compensatory neural effort to sustain performance, whereas Nature facilitated efficient, low-effort processing through stable neural regulation.

**Figure 10.**
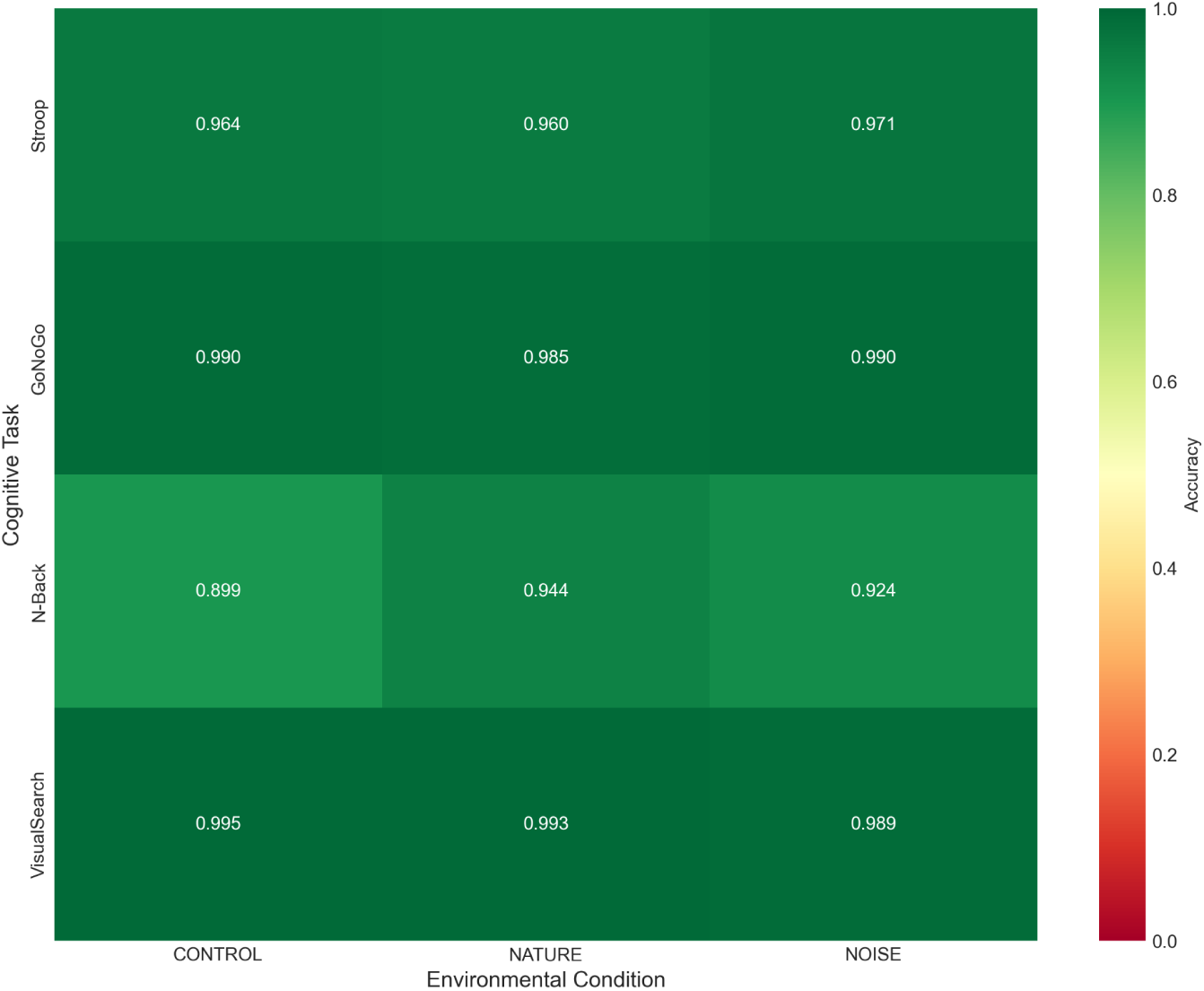
Cognitive Task Performance Across Environmental Conditions. Heatmap showing mean accuracy for four cognitive tasks (Stroop, GoNoGo, NBack, VisualSearch) across Control, Nature, and Noise conditions. Color indicates accuracy (0.0 = *red*, 1.0 = *darkgreen*).

### 3.5 Environmental Neurological Risk (ENR) Index

ENR Index scores across participants (*n* = 12) ranged from 9.5 to 18.0 (*M* = 14.8, *SD* = 2.0), with 91.7% (*n* = 11) classified as good health (ENR ≥ 14) and one participant (8.3%) as moderate health (7.5 ≤ ENR *<* 14; Figure11A). No participants fell below the low health threshold (ENR *<* 7.5). The distribution provided sufficient variance to explore individual differences in environmental sensitivity, with participant P006 (ENR = 9.5) exhibiting the lowest score. To test whether baseline health status moderated neural responses to environmental context, baseline-corrected condition effects were computed by subtracting resting-state activity from task-related neural activity (task – baseline), and then comparing Nature–Control differences as indices of restorative neural response. All correlations were adjusted for multiple comparisons using FDR correction. Across nonlinear EEG features, the strongest association with ENR Index emerged for Hjorth complexity during Nature exposure (Figure11B–D). Higher ENR scores were associated with greater increases in baseline-corrected Hjorth complexity under Nature relative to Control (*r* = 0.615, uncorrected *p* = 0.033), representing the largest effect size identified across 36 correlations tested. After FDR correction, however, this effect did not reach significance (*p* = 0.997). The positive trend suggests that individuals with better overall health may exhibit stronger neural plasticity and adaptive cortical reorganization in response to restorative environments. Other nonlinear features showed weaker or nonsignificant relationships (all FDR-corrected p = 0.997):

- DFA *α* (long-range temporal correlations): *r* = 0.264
- Sample entropy (signal irregularity): *r* = 0.07
- Spectral entropy (frequency irregularity): *r* = −0.08
- Hurst exponent and Hjorth mobility: both *r* ≈ 0.2 − 0.3

Among the top six nonlinear features examined, Hjorth complexity displayed the largest positive relationship with Nature-related restoration (r≈ 0.6), followed by smaller positive trends for DFA *α* and Hurst exponent, whereas entropy-based measures clustered near zero (Figure 11C-D). None of the spectral power bands (*δ*, *θ*, *α*, *β*, *γ*) correlated significantly with Nature restoration effects (all FDR-corrected *p* = 0.997). In contrast, ENR Index correlations with Noise-induced stress responses were weak or near zero across all nonlinear features (e.g., spectral entropy r≈ −0.2, sample entropy r≈ −0.2, Hjorth mobility r ≈ −0.3, all FDR-corrected p = 0.997). Comparison of Nature versus Noise moderation effects revealed an asymmetry: features showing the strongest Nature-related associations (Hjorth complexity, DFA *α*, Hurst exponent) showed negligible correlations with Noise responses (|r| < 0.35). This suggests that restorative benefits of Nature may depend more strongly on baseline health status than stress-related costs of Noise. Individual analyses supported this interpretation: participant P006 (ENR = 9.5) showed one of the weakest Nature-induced increases in Hjorth complexity (Figure 11B), consistent with the overall positive trend. In summary, ENR Index analyses using baseline-corrected neural activity revealed exploratory evidence that baseline health status may moderate Nature-related restorative neural responses, particularly through Hjorth complexity (r = 0.615, uncorrected p = 0.033). None of the tested relationships survived FDR correction (all p = 0.997), indicating that these effects should be interpreted as preliminary. Nonetheless, the direction and magnitude of the Hjorth complexity effect (37.8 % variance explained) suggest that neural signal complexity, reflecting cortical flexibility and adaptive capacity, may capture early signatures of health-dependent environmental restoration. This aligns with the concept of neural “criticality,” in which healthy brains maintain an optimal balance between stability and variability, supporting efficient adaptation to restorative environmental contexts.

**Figure 11.**
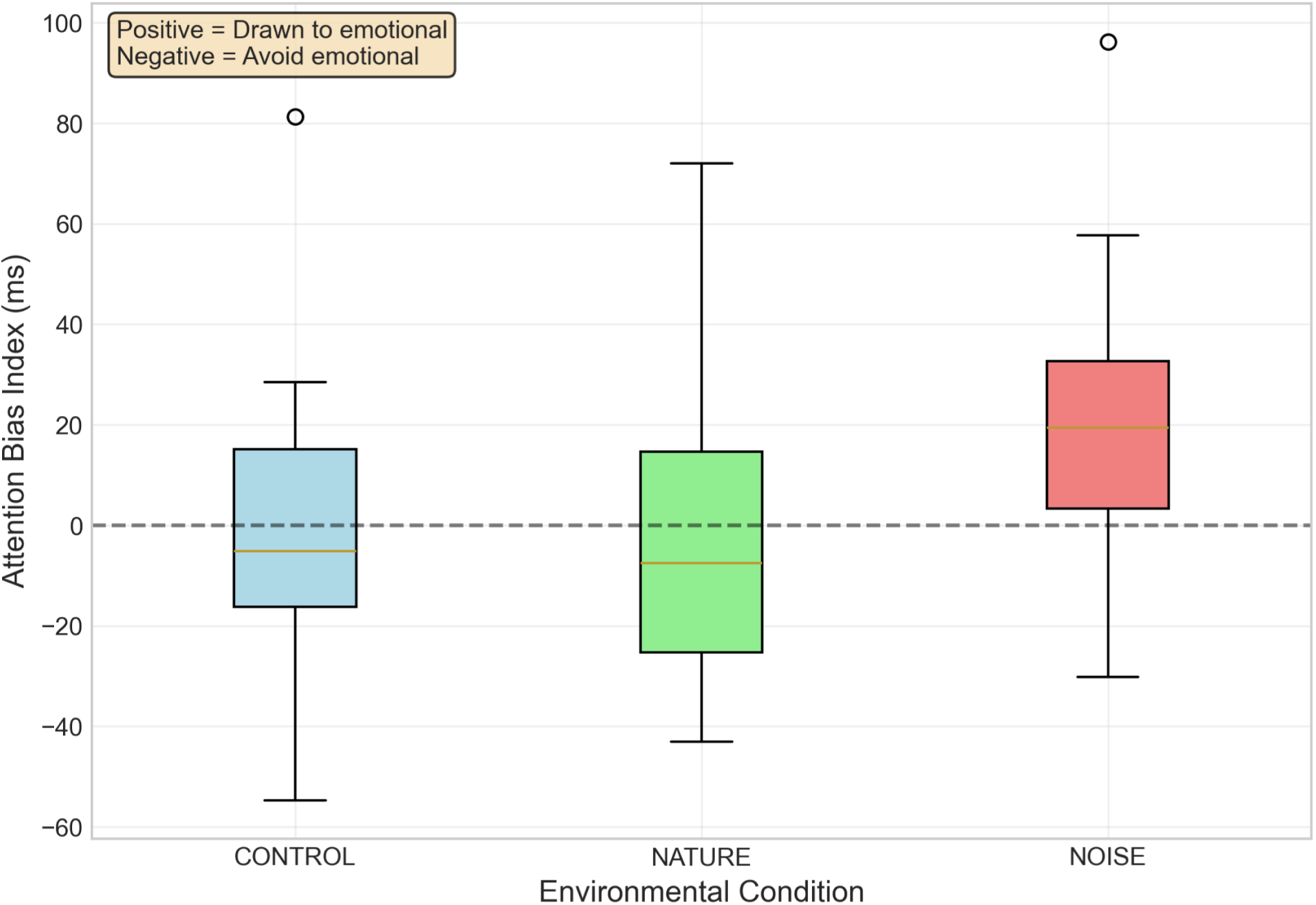
Attention Bias by Environmental Condition. Box plot showing Attention Bias Index (ms) across Control, Nature, and Noise conditions. Positive values indicate attention drawn to emotional stimuli; negative values indicate avoidance. Control: median = −2 ms (neutral). Nature: median = −8 ms (slight avoidance). Noise: median = 19 ms (drawn to emotional), with entire IQR above zero.

**Figure 12.**
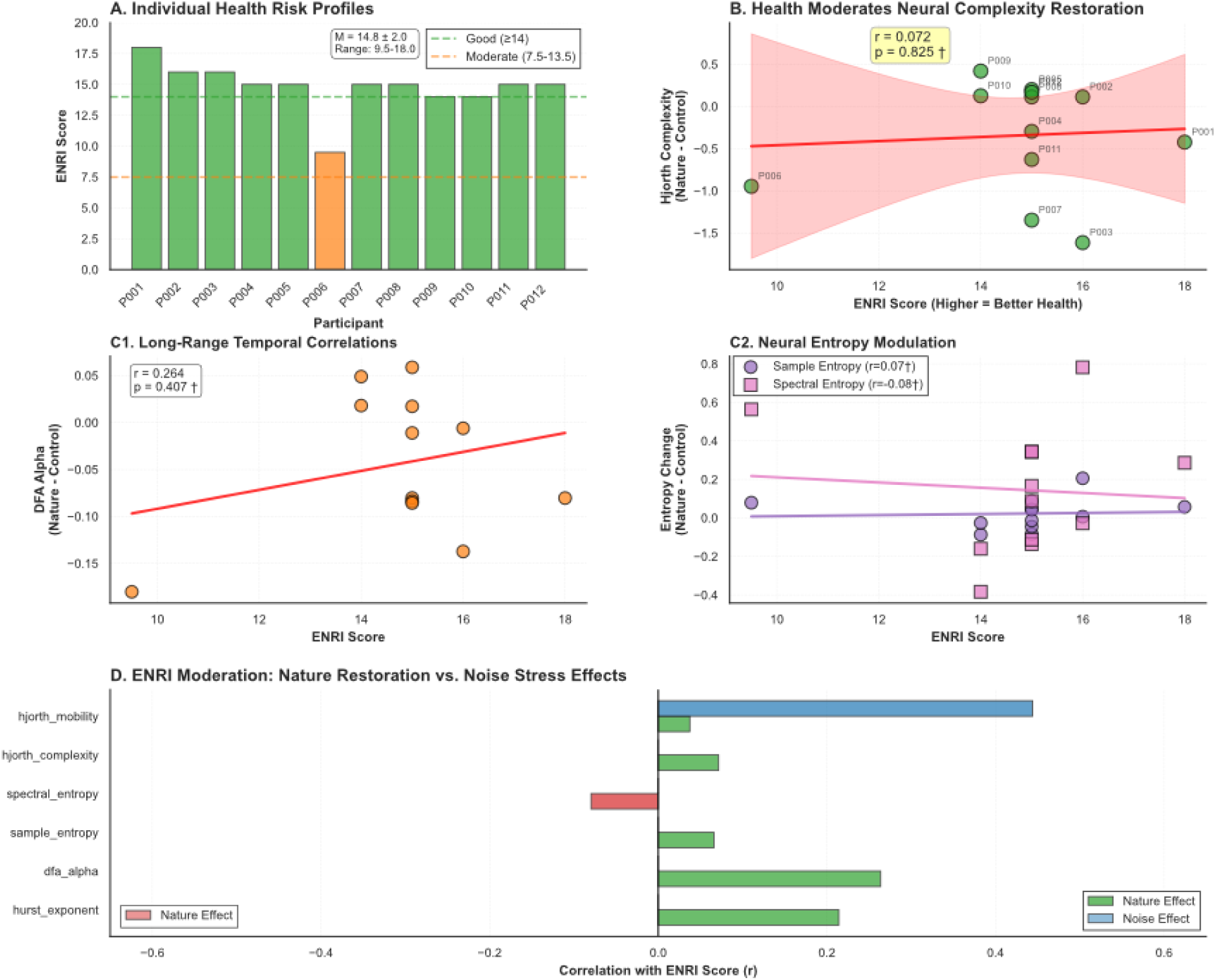
ENR Index Moderation of Environmental Neural Responses. (A) Bar chart of each participant’s ENR Index score, color-coded by risk category (Green = Good 20-14, Orange = Moderate 7.5-13.5, Red = Low <7.5), with threshold lines and summary statistics. (B) Scatter plot showing correlation between ENR Index and Hjorth Complexity Nature restoration effect. Red regression line with 95% CI, participant labels, and statistics box (r and p-value). (C1) DFA Alpha correlation with ENR Index (orange points, marginal trend), (C2) Sample Entropy (purple circles) and Spectral Entropy (pink squares) correlations with ENR Index. (D) Horizontal bar chart comparing Nature restoration effects versus Noise stress effects across top 6 neural features.

## 4 Discussion

The present study demonstrates that everyday auditory environments, specifically natural sound-scapes versus urban noise, produce dissociable effects on cortical dynamics, cognitive performance, and emotional attention. Using multimodal EEG analyses spanning spectral, ratio-based, and nonlinear domains, we show that environmental sound systematically reorganizes large-scale neural activity even in the absence of overt behavioral impairment. At a systems level, natural soundscapes promote efficient, low-effort neural states, whereas urban noise elicits compensatory, strain-related activation consistent with stress-laden processing. These results bridge environmental neuroscience, cognitive electrophysiology, and health science, and align with theoretical frameworks on restoration, predictive processing, and allostatic regulation. (Kaplan, 1995; Berman et al., 2008; Friston, 2010; McEwen and Gianaros, 2011)

### 4.0.1 Environmental context shapes cortical efficiency

Across analyses, Nature and Noise exposures evoked distinct large-scale neural profiles. Nature reflected a state of relaxed alertness that optimally balances arousal and control central to Attention Restoration Theory (ART). In contrast, Noise heightened cognitive load and disrupted attentional regulation. This dissociation supports theories of neural efficiency and environmental restoration, whereby restorative environments facilitate stable and energetically economical brain states, while high-load environments impose compensatory resource demands. (Kaplan, 1995; Berman et al., 2008) Mechanistically, predictive coding suggests that coherent, low-entropy input (e.g., nature) stabilizes internal models and reduces prediction error, lowering metabolic demand; conversely, spectrally and temporally unpredictable noiseincreases error signals and resource consumption, pushing networks toward less efficient regimes. (Friston, 2010). Complementarily, allostatic load theory predicts that maintaining performance under stress incurs physiological “wear-and-tear,” matching our observation of stable behaviour with elevated neural effort under noise. (McEwen, 2007; McEwen and Gianaros, 2011).

### 4.0.2 Neural-behavioral dissociation and compensatory activation

Despite pronounced neural modulation, behavioral performance remained near ceiling for most tasks. Only the working-memory (NBack) task exhibited some measurable differences, with accuracy improving under Nature relative to Noise and Control. The coexistence of stable behavior with divergent neural patterns indicates a neural-behavioral dissociation, whereby cortical systems expend additional resources under Noise to maintain equivalent output. This compensatory activation mirrors predictions from cognitive energetics and allostatic load models, suggesting that noise exposure pushes neural systems toward less efficient operating regimes, achieving performance at the cost of increased effort and reduced regulatory flexibility. Such dissociations caution against relying solely on behaviour to index cognitive costs of environments, since neurophysiological indices reveal latent strain that may matter for fatigue and well-being over longer horizons (McEwen and Gianaros, 2011).

### 4.0.3 Complexity-based signatures of adaptive restoration

Exploratory analyses of the Environmental and Neurological Risk (ENR) Index suggests a potential moderating role of baseline health in shaping neural responses to restorative contexts. Although the Hjorth complexity-ENR relationship did not survive correction for multiple comparisons, the positive association (r = 0.615) suggests that individuals with better overall health exhibit greater increases in neural signal complexity during Nature exposure. Hjorth complexity measures are closely linked to neural criticality, the dynamic balance between stability and variability that optimizes information processing. Higher complexity under Nature thus may show a capacity for adaptive reorganization and efficient environmental integration, while lower complexity (as seen in the participant with the lowest ENR score) may reflect reduced neural flexibility. These findings suggest that restorative neural plasticity may depend on an individual’s baseline resilience, a hypothesis consistent with emerging work on health-related variability in brain-environment coupling (Marcantoni et al., 2023).

### 4.0.4 Environmental modulation of attention and emotion

The behavioral data complement this neurophysiological picture. Nature soundscapes preserved accuracy and attentional balance across all tasks, whereas Noise selectively reduced working-memory precision and increased orienting toward emotional cues in the DotProbe task. This shift toward emotionally salient stimuli under Noise aligns with limbic-attentional coupling models, in which unpredictable sensory environments bias attention toward threat-related signals. These results indicate that environmental sound not only shapes cortical efficiency but also influences the balance between cognitive control and emotional reactivity (Bar-Haim et al., 2007).

### 4.0.5 Mechanistic and translational implications

Taken together, the results advance a mechanistic model in which Nature supports efficient predictive processing by stabilizing cortical networks through reducing sensory uncertainty and energetic demand. By contrast, Noise increases prediction error and metabolic load, driving compensatory activation and reduced signal complexity. These findings resonate with frameworks from predictive coding and allostatic regulation, in which natural auditory environments provide coherent, low-entropy sensory input that reinforces internal model stability, whereas noise imposes unpredictable perturbations that challenge homeostatic balance. This means that the data supports biophilic interventions in workplaces (Bar-Haim et al., 2007) and designing soundscapes should therefore consider both behavioral endpoints and neural economy (Berman et al., 2008). From an applied perspective, the combination of *α/β* ratio, Engagement Index, which has since been validated across vigilance and learning contexts, and Hjorth complexity, may serve as sensitive indicators of cortical efficiency and adaptive capacity. Together they represent quantitative neural markers for further environmental health research and human-health applications (Pope et al., 1995; Allen et al., 2004; Coan and Allen, 2004; Puma et al., 2018). Baseline reliability analyses further revealed that Nature soundscapes reduced the temporal stability of spontaneous EEG activity relative to Control and Noise, a pattern consistent with enhanced neural flexibility and exploratory dynamics in restorative contexts. This interpretation aligns with the view that natural environments promote greater moment-to-moment variability, which is an adaptive feature of low-arousal and self-organized cortical states, rather than instability necessarily.

### 4.0.6 Methodological considerations

Two analytic choices deserve emphasis. First, baseline-correction (task-baseline) isolates environment-specific task modulation from trait/state baseline differences, increasing sensitivity in environmental EEG. Second, mixed-effects modelling is well-matched to repeated-measures EEG, alongside FDR control (Benjamini-Hochberg), where it guards against inflated false positives. Notably, cluster-based permutation tests were null here, reminding that non-clustered but systematic effects can arise when manipulations are moderate and distributed, an expected scenario in ecological exposures (Benjamini and Hochberg, 1995; Maris and Oostenveld, 2007).

### 4.0.7 Limitations and future directions

The sample size (n = 12) limits generalizability and precludes definitive inference regarding interindividual moderation. Moreover, the strictly auditory manipulation captures only one facet of real-world exposure; future work should extend these paradigms to multimodal and longitudinal designs, integrating physiological and ecological measures (e.g., heart-rate variability, pollution exposure, and heat exposure). Replication in larger, demographically diverse cohorts will be critical to confirm whether complexity-based neural indices reliably track environmental resilience (Loo and Makeig, 2012; Arns et al., 2013).

### 4.0.8 Conclusions

Together, these findings identify distinct electrophysiological signatures of restorative versus stress-inducing environments. Natural soundscapes promote engaged relaxation and adaptive neural complexity, enabling efficient cognitive functioning with minimal strain. Urban noise elicits compensatory high-effort states, marked by greater slow-wave synchronization and reduced flexibility. Despite stable overt performance, the underlying neural economy diverges sharply, revealing that the brain’s response to environmental context is both measurable and mechanistically meaningful. By integrating multiscale EEG metrics with behavioral and health data, this study contributes to a growing body of evidence positioning the environment as a modulator of cognitive loads and neural organization. It supports a view of the brain as an environmentally embedded predictive system, constantly adjusting its internal dynamics to the sensory regularities or unpredictabilities of the world around it (Friston, 2010).

## Supplementary Information

### 4.1 Supplementary Figure 1

**Sup. Fig 1.**
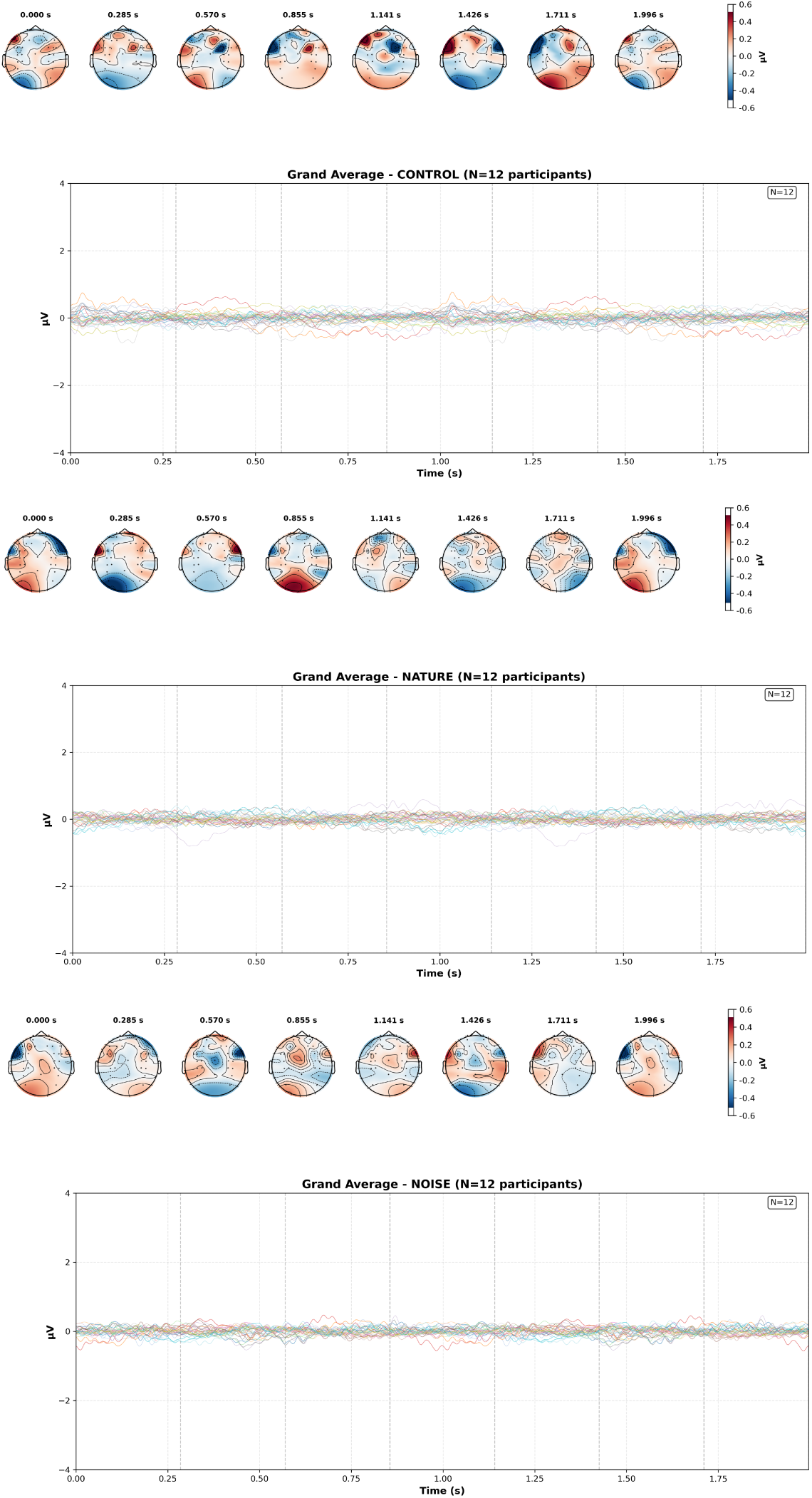
Grand Average Evoked Responses Across All Participants by Environmental Condition Grand average evoked responses (ERPs) computed across all 12 participants (N=12) for each of the three auditory conditions: Control, Nature, and Noise. Each panel follows the same format: the upper section displays eight topographic maps showing the spatial distribution of electrical potential across the scalp at evenly spaced time points throughout the 2-second epoch. The lower section shows the time course of the grand average evoked response across all 32 EEG channels, with each channel represented by a colored trace. The color scale for topographic maps ranges from −0.6 ÂţV (dark blue, negative potentials) to +0.6 ÂţV (dark red, positive potentials), with white/light blue indicating values near baseline. The time course plots display voltage (ÂţV) on the y-axis (−4 to +4 ÂţV) and time (seconds) on the x-axis (0.00 to 1.75 s), with vertical dashed lines indicating the time points corresponding to the topographic maps displayed above.

**Sup. Fig 2.**
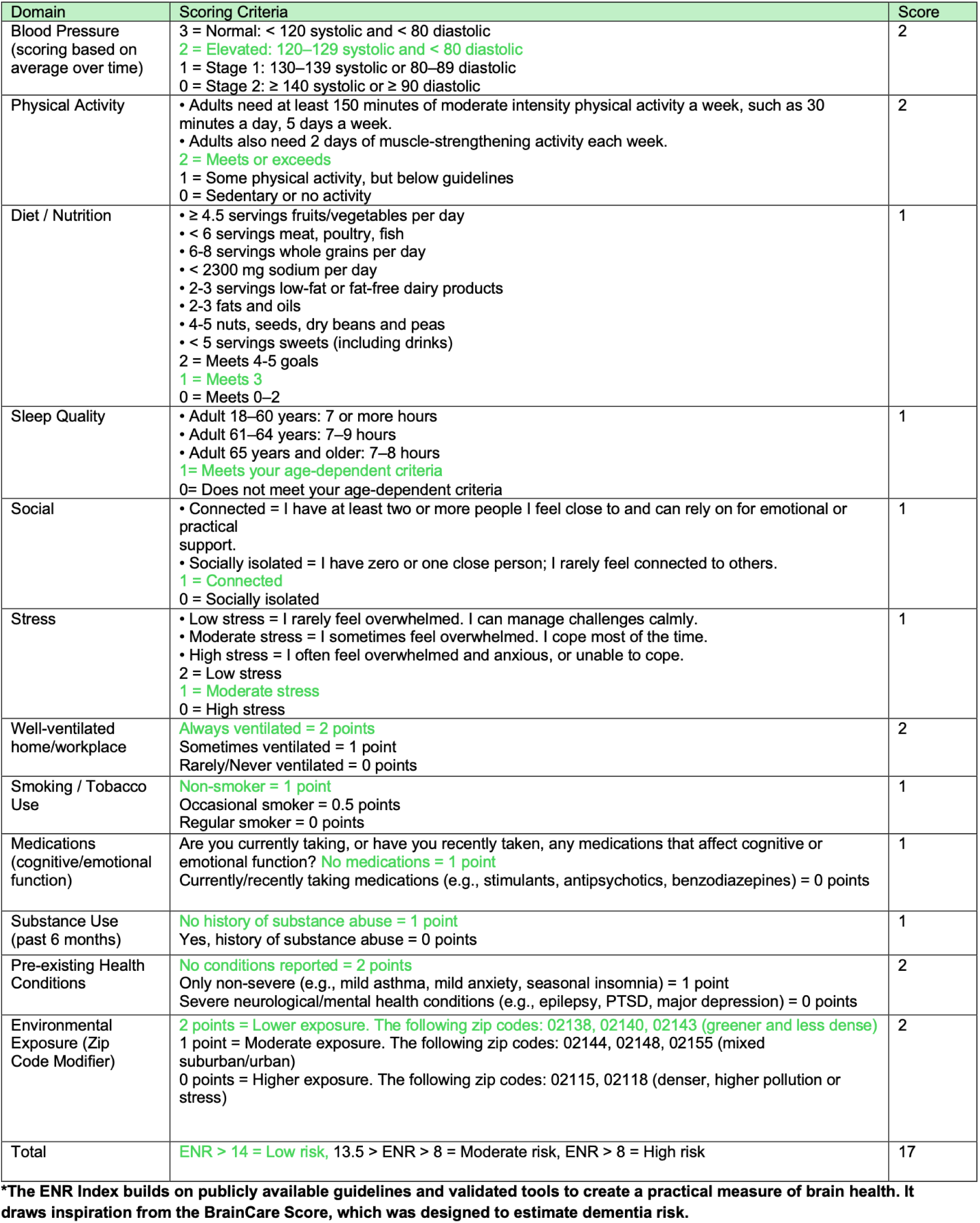
Environmental Neurological Risk (ENR) Index Illustrative example (shown in green) of how an individual’s ENR Index is computed across domains. Each category-covering physiological, lifestyle, psychological, and environmental factors-is scored based on publicly available guidelines and validated tools. The summed ENR Index provides an overall brain health risk classification

## Acknowledgments

The author thanks her husband, Dr. Federico Claudi, for his insightful comments and scientific feedback on earlier versions of this manuscript. The author also thanks her undergraduate research assistants, Anita Avdiu and Andi Buqa, for their help in refining participant survey questions.

## Author Contributions

E.M. conceived and designed the study, conducted all data collection and analyses, and wrote the manuscript. Large language models were used to assist in language clarity.

## Data and Code Availability

The datasets and analysis code supporting this study are part of ongoing development for a forthcoming application and cannot be made publicly available at this stage. Processed, de-identified data and analysis scripts will be shared upon reasonable request once intellectual-property protection and app deployment are complete.

